# Theta activity in the RSC anchors space to the body cardinal axes

**DOI:** 10.64898/2025.12.02.691796

**Authors:** Clément Naveilhan, Stephen Ramanoël

**Author notes:** CA, Corresponding Author: Clément Naveilhan & Stephen Ramanoël.

## Abstract

Understanding human navigation in ecological, freely moving conditions requires uncovering how the brain anchors directional representations to the body’s orientation. Using high-density mobile electroencephalography and immersive virtual reality during goal-directed whole-body rotations, we found that theta bursts reconstructed in the retrosplenial complex (RSC) encode both acceleration and alignment with the body’s principal axes. Crucially, this body-axis–anchored neural signal emerged only during goal-directed rotations, and its strength correlated with individual navigation performance, suggesting an adaptive mechanism which provides a stable egocentric scaffold for orientation. These results provide compelling evidence for a self-motion-gated, body-centered reference frame that supports efficient navigation, and bridge the gap between static neuroimaging findings in humans and rodent research on RSC geometry codes. Overall, our findings advance an embodied, mechanistic account of human navigation, opening new avenues for investigating brain dynamics in naturalistic, movement-rich settings using non-invasive neural recordings.

## Introduction

Spatial navigation arises from the intricate coordination of multiple cognitive and sensorimotor processes. At the core of this complexity lies path integration, the ability to update one’s position when deprived of external information by integrating self-motion cues (*e*.*g*., vestibular and proprioceptive). Relying solely on such cues, humans can accurately track their location to reach an intended destination (McNaughton et al., 2006). However, navigating under such conditions deprived of any external cues is inherently difficult, and errors accumulate rapidly (Stangl et al., 2020). To support efficient navigation in this context, spatial representations can be organized around bodily reference axes (*i*.*e*., cardinal body axes) that provide a stable scaffold for orientation and memory when environmental cues are sparse or unreliable. Despite its importance, how the human brain implements this embodied spatial anchoring during active navigation remains unclear.

Only a handful of recent studies have begun to address this question, implicating the retrosplenial complex (RSC) as a central locus of these egocentric processes. Located at the interface of visual and memory circuits, this medial parietal region acts as an integrative hub of multimodal signals and serves as a neural compass (Alexander et al., 2023; Dutriaux et al., 2024; Lu et al., 2025; Stacho & Manahan-Vaughan, 2022). For example, Dutriaux *et al*. (2024) found that the RSC carries egocentric codes for both one’s facing direction and the direction of the goal. Similarly, Lu *et al*. (2025) combined immersive virtual-navigation paradigms with functional magnetic resonance imaging (fMRI) and reported that the RSC exhibits stable egocentric directional tuning relative to the principal axis of the environment. Activation patterns in the RSC varied consistently with the subject’s facing direction and with the goal direction when defined relative to the body, rather than only relative to the external map. These findings suggest that the RSC can flexibly support orientation and goal□directed navigation by anchoring directional codes to a body-centered egocentric frame, rather than a purely allocentric reference frame (Alexander et al., 2023).

However, these studies relied on static, supine MRI paradigms, thereby disregarding vestibular and proprioceptive self-motion cues that are fundamental for accurate spatial navigation (Stangl et al., 2023; Taube et al., 2013). From this perspective, in a recent study our research group leveraged advances in virtual reality and mobile electroencephalography (mobile EEG) to extend the known role of the RSC in naturalistic human navigation (Naveilhan et al., 2025). We reported that theta activity in the RSC plays a dual role, supporting both landmark-based spatial recalibration and the encoding of self-motion signals at rotation onset. These findings position RSC theta oscillations as a key mechanism for spatial anchoring, aligning egocentric motion cues with the allocentric structure of the environment. Yet whether these oscillations also implement the body-axis anchoring suggested by a few behavioral and recent fMRI studies remains unknown. Understanding this link is essential to connect behavioral evidence of body-axis anchoring with the neural computations in the RSC that support real-world spatial orientation.

To this end, we recorded EEGs from 40 participants who physically rotated to indicate the direction of a target in immersive virtual reality. Analysis of dipoles reconstructed around the RSC revealed that bursts of theta activity selectively encoded alignment with the body’s principal axes. This body-axis reference frame signal arose only during goal-directed rotation and scaled with behavioral accuracy, indicating an adaptive mechanism that provides a stable egocentric scaffold for orientation.

## Results

### RSC theta activity supports anchoring of space around the left-right axis

Our first analysis examined whether theta-episode onsets were preferentially triggered when participants crossed the left–right body axis (90°/270°; **Fig. 1A–B**). We extracted the onsets of theta episodes, which were binned in 3° sectors (excluding 0–3° and 357–360° as participants were immobile for most of the time in this position facing the 2D map). We applied a Poisson generalized linear mixed-effects model (GLME, log link) comprising circular predictors (cos θ, sin θ) and Gaussian anchor regressors (σ = 30°) centered at 90° and 270° (**Fig. 1C**). Adding the anchor terms markedly improved model fit over a circular-only model (Likelihood-ratio χ^2^ = 1269.06, *p* < .001). Both anchors were significant: 90° (*β* = 0.946 ± 0.044, z = 21.45, *p* < .001; Incidence Rate Ratio = 2.57, 95% CI [2.36, 2.81]) and 270° (*β* = 1.090 ± 0.044, z = 24.55, *p* < .001; IRR = 2.97, 95% CI [2.73, 3.25]). These effects indicate an increase of approximately 2.5- to 3-fold in theta-episode incidence near the left– right body axis relative to the baseline circular pattern, demonstrating a marked concentration of theta bursts around this lateral body axis. Importantly, this pattern was not explained by movement-related variables such as acceleration (**Fig. 1D**) or speed (**Fig. 1E**). Although acceleration reliably triggered theta bursts, consistent with our previous findings (Naveilhan et al., 2025), this effect was restricted to small angles during initiation of the rotation and did not account for the axis-specific anchoring. Together, these results provide compelling evidence that theta burst onsets are preferentially anchored to the cardinal left–right axis.

**Figure 1.**
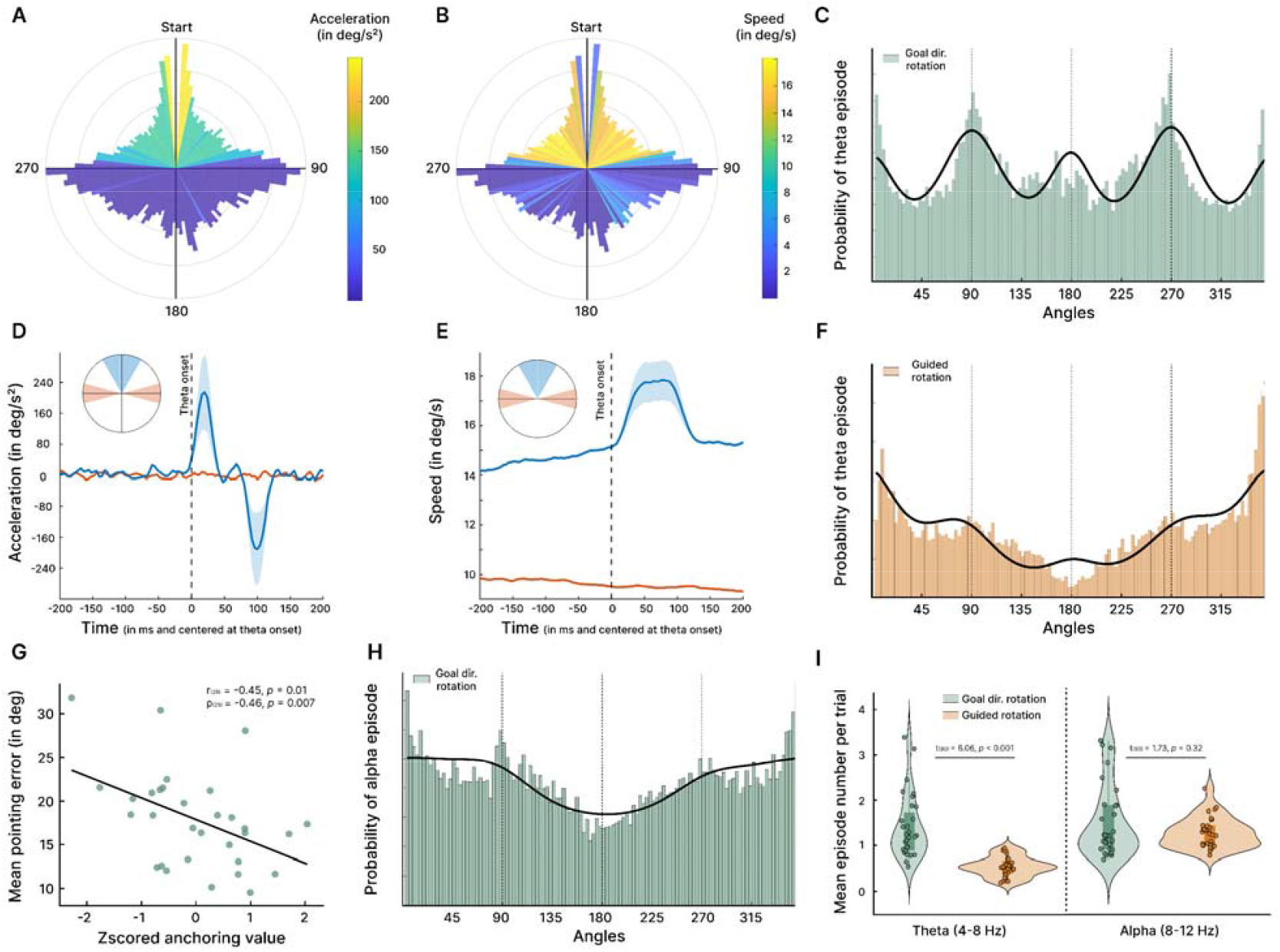
Theta activity specifically anchors space around the left-right body axis, only during goal directed rotation and it correlates with pointing performance. **A**. Polar histogram representing the probability of theta episodes pooled across all subjects and binned into 3° intervals. Color indicates the mean acceleration for each angular bin. **B**. Same as in A, but color-coded by mean speed. **C**. Linear histogram showing the probability of theta episodes across angular bins (equivalent to A, but unwrapped), with a Poisson generalized linear mixed-effects model (GLME, log link) fit. Predictors included circular components (cos□θ, sin□θ) and Gaussian-shaped regressors (σ□=□30°) centered at 90° and 270°. **D**. Time-locked average acceleration traces centered on theta episode onset, separated for small-angle episodes near 0° and episodes near the 90°/270° axis. **E**. Same as D, but for speed. **F**. Histogram of theta episode probability during guided rotations, with a similar GLME model applied as in C. **G**. Correlation between individual mean pointing error and the z-scored anchoring strength, derived from the average beta weights at 90° and 270° in the GLME. **H**. Angular histogram of alpha episode probability during goal-directed rotations. **I**. Comparison of the number of detected episodes in the theta (4–8□Hz) and alpha (8–12□Hz) bands during goal-directed versus guided rotations.

**Figure 2.**
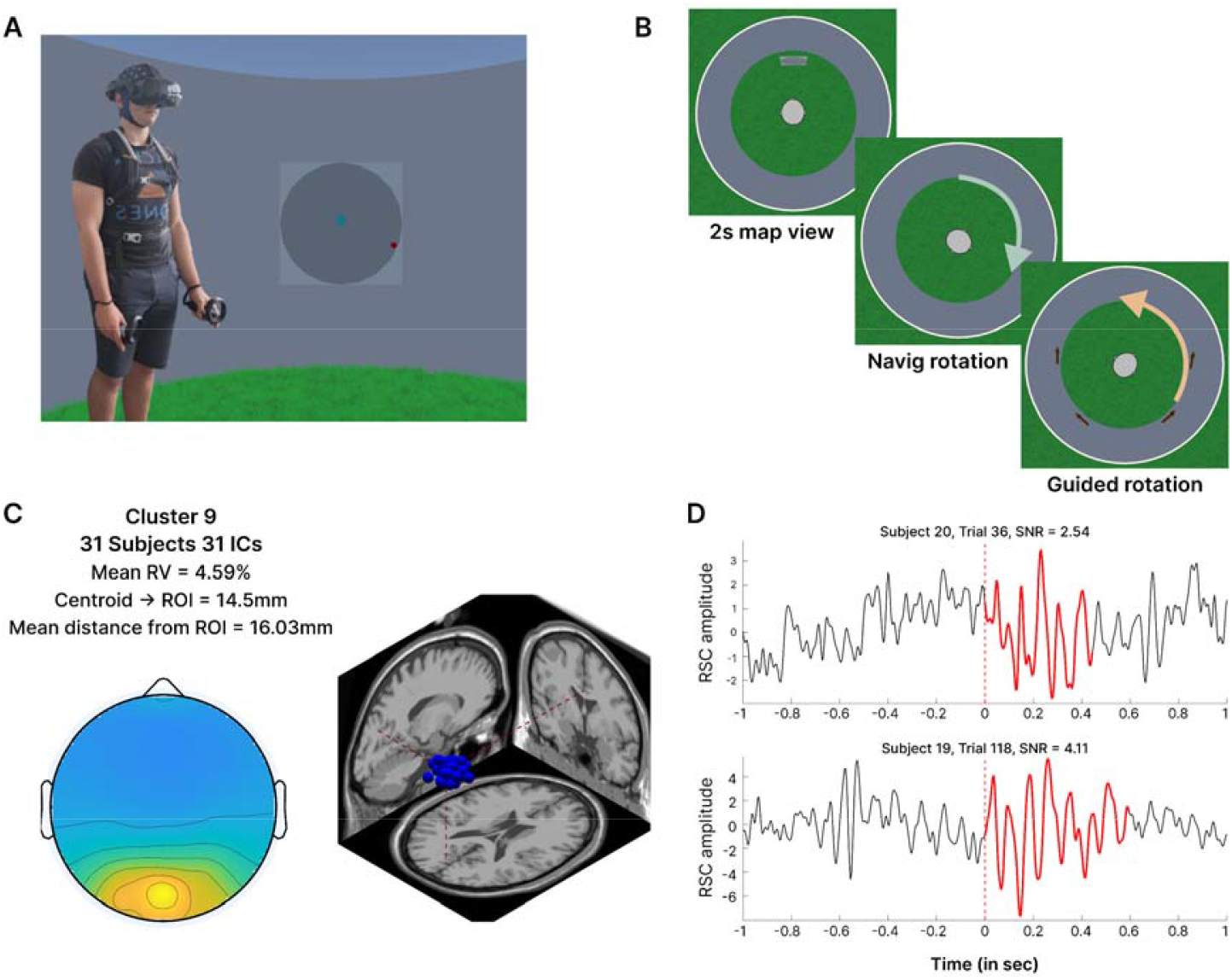
Presentation of the protocol and the EEG analysis. **A**. Experimental setup. Participants were equipped with a backpack-mounted recording system and wore an immersive head-mounted display (HTC Vive Focus 3), along with a 64-channel ANT Neuro Waveguard EEG cap. During free rotation in the virtual environment, neural and motion data were recorded and synchronized with millisecond precision using the Lab Streaming Layer (LSL). **B**. Task protocol. Each trial began with the presentation of a map displaying a red dot indicating the target direction. After 2 seconds, the map disappeared, and participants were free to rotate and stop when they believed they were facing the target direction. Four directional arrows then appeared, to guide them back to the initial orientation. Once they returned to the 0° starting position, the map was re-displayed, initiating the next trial. **C**. EEG source analysis. Following preprocessing, source-level dipoles were reconstructed from scalp EEG data and clustered across participants. After 5,000 iterations, the optimal clustering solution was selected, retaining data from 31 participants. In cases where multiple dipoles from a single subject were included, the one closest to the region of interest (ROI) was retained, yielding a mean dipole–ROI distance of 16.03 mm. **D**. Example of detected EEG bursts. Two selected theta bursts are shown, occurring near 90° angular transitions (0 on the x-axis indicates the moment participants passed through 90°). For visualization, the signal was bandpass filtered between 2–30□Hz.

### Body-axes anchoring is specific to the goal-directed rotation phase

We then conducted the same analysis but on data obtained during the guided rotation phase, in which participants followed the directional arrows to return to the starting orientation (**Fig. 1F**). This analysis revealed no significant anchoring near the left-right body axis: 90° (z = 1.629, *p* = .103) and 270° (z = 1.022, *p* = .307). However, during these guided rotations, theta activity increased markedly as participants approached the starting orientation that corresponded to their principal body axis throughout the task. This effect was captured by a Gaussian kernel centered at 0°, which improved fit over a circular-only model (likelihood-ratio χ^2^ = 319.96, *p* < .001). In this Poisson GLME, the 0° term was positive and precise (*β* = 0.621 ± 0.056, z = 11.25, *p* < .001), corresponding to an ∼86% higher incidence of theta onsets near 0° (IRR = 1.861, 95% CI [1.670, 2.074]). To identify the preferred theta-firing direction, we folded angles into the 0–180° range using 1° bins and applied linear statistics. This revealed a pronounced peak at 5.5° (5°-6° bin), with theta-episode counts increasing 2.71-fold relative to a uniform null (IRR = 2.706, 95% CI = [2.357, 3.105], *p* _20000 permutations_ < .001). The observed increase in theta activity near the starting position complements the previous observation of increased theta around the cardinal left–right axis. This pattern supports a four-fold symmetry model of RSC spatial coding, where the alignment of the principal axis with the starting orientation anchors the construction of the egocentric left–right reference frame, as reported at the behavioral level (He & McNamara, 2018). However, fully establishing such a four-fold pattern would require assessing the 180° axis, which was not possible in our current study because participants rarely crossed 180° (9.7 ± 7.2 trials per subject, vs. 171.09 ± 17.3 for the left–right axis), precluding a reliable analysis of that direction. Despite this limitation, we still observed a decrease in pointing error at 180°, mirroring the effect seen along the left–right axis, suggesting that a similar directional stabilization may also operate at this orientation (Saulay-Carret et al., 2025).

### Neural signature of body anchoring correlates with pointing accuracy and is specific to theta activity

We then tested whether pointing performance (angular error) was related to the strength of the anchoring. To this end, firstly we modeled the pointing performance in the GLME, but it did not improve the model compared to the one without the interaction (likelihood-ratio χ^2^ = 0.49, *p* = .91). This was confirmed by the absence of a main effect of performance (*z* = 0.23, *p* = 0.82) and by the lack of interaction between performance and anchoring for both 90° (*z* = – 0.10, *p* = .92) and 270° (*z* = –0.63, *p* = .53). However, when considering the error at the subject level, we reported a negative correlation between the strength of anchoring (as expressed in z-scored value for averaged *β* at 90° and 270°) and the error in pointing of the subjects (Pearson correlation: r = -0.453, *p* = .010 and Spearman correlation : ρ = -0.464, *p* = .007; **Fig. 1G**). These results suggest that participants who performed best on the pointing task also showed the strongest body-axis anchoring via theta activity, highlighting the adaptive nature of this egocentric mechanism in an environment deprived of any other cues.

In a control analysis, we assessed the specificity of the theta effect by fitting an equivalent GLME to RSC alpha activity (8–12 Hz; **Fig. 1H**), given prior evidence implicating this alpha band in the computation of heading changes during spatial orientation (Lin et al., 2015). Here we reported an absence of anchoring for both 90° (z = 1.011, *p* = .312) and 270° (z = 0.513, *p* = .608), highlighting that the effect is specific to theta activity. We also compared the average number of episodes per subject, per trial for both theta and alpha activity using linear mixed models (**Fig. 1I**). Our results reported a main interaction between the frequency and the rotation (F_(1,90)_ = 9.45, *p* = .002), and while we reported an increased number of theta episodes for the goal-directed compared to the guided phase (t_(90)_ = 6.06, *p* < .001), we reported no effect for the alpha band (t_(90)_ = 1.71, *p* = .32). We observed a similar absence of anchoring for delta (2-4 Hz, all *p* > .45) and beta band activity (12-30 Hz, all *p* > .43). Taken together, these results strengthen the specificity of theta activity for anchoring space through this cardinal body axis, and only when participants were actively aiming to reach an intended spatial orientation.

## Discussion

In this mobile EEG study, we investigated the neurocognitive mechanisms underlying the egocentric anchoring of space around cardinal body axes during orientation. We found that during goal-directed rotations, the RSC displayed an increased number of theta episodes when participants crossed these axes, despite the absence of any external visual cues indicating this orientation. Critically, this pattern was only observed during these goal-directed rotations, and not when participants were guided to perform the same rotation to begin a new trial. These results suggest that theta activity in the RSC reflects an egocentric four-fold symmetry pattern highlighting an adaptive anchoring of space around the main body axes to support spatial orientation.

Taking advantage of a new dataset, we firstly confirmed previous findings that theta activity was related to the encoding of vestibular information during rotation (Gramann et al., 2021; Naveilhan et al., 2025). In the present study, using a burst detection approach, we confirmed that the acceleration phase of the rotation was associated with the onset of theta episodes, as previously proposed in animal models (Hennestad et al., 2021; Kropff et al., 2021). These theta bursts could reflect the integration of vestibular acceleration signals to update head-direction estimates in real time, consistent with frameworks linking vestibular drive to the rapid recalibration of spatial orientation (Laurens & Angelaki, 2017; Yoder & Taube, 2014).

Our results further extend the role of theta activity, showing that it not only occurred upon rotation onset, but also increased when participants faced the left-right body axis. This is particularly notable as participants could not use any external information that could trigger this activity. This pattern suggests that RSC theta activity supports an internally generated spatial anchor aligned with the cardinal body axes, echoing behavioral evidence that memory is facilitated for items aligned with an intrinsic layout axis (Mou & McNamara, 2002). Convergently, He and McNamara (2018) found that a body-centered reference frame (front– back and left–right) governs performance in spatial updating. They showed that an initial heading can anchor these axes, as we similarly found with increased theta activity close to the starting position. We also previously observed a facilitating effect for the angle aligned with the cardinal body axes in a previous study (Saulay-Carret et al., 2025) and the present neural results offer a possible explanation for these findings. In our task, participants could rely exclusively on self-motion cues, thus relying on egocentric strategies. In these conditions, people tend to use the body axes as a reference system, with a four-fold symmetry allowing the comparison and adjustment of their actual position relative to these main axes (Sigismondi et al., 2024; Wagner et al., 2023).

In line with this account, using fMRI methods Bécu *et al*. (2025) observed that human RSC expresses both a body-axis–linked egocentric code and an allocentric, multi-axis “clover” code. In their results, RSC response was maximal when remembered targets lay behind the avatar, and independently exhibited four-fold directional tuning anchored to the virtual arena’s dominant axes. Our results extend this picture by showing that a similar four-fold symmetry pattern emerges when participants physically perform the rotations, even in the absence of landmarks, with anchoring around the principal body axes. This finding underscores the flexibility of the RSC in dynamically tracking and integrating the most relevant information for navigation, whether derived from external or self-motion signals (Alexander & Nitz, 2015).

Our findings provide the first evidence of this egocentric anchoring in freely moving humans, but it is important to highlight that they closely align with animal data. In rodents, RSC neurons exhibit multidirectional tuning reflecting environmental geometry, including clear four-fold (90°) periodicity in square arenas (Zhang et al., 2022). Extending this, LaChance and Hasselmo (2024) showed that the RSC encodes local geometric axes with stronger four-fold symmetry, while the postrhinal cortex captures global boundary layout. Finally, Park *et al*. (2024) identified “vertex cells” in granular RSC that fire at environmental corners at specific egocentric bearings and distances, and were enhanced during goal-directed navigation, paralleling the selectivity we observed. Together, these studies demonstrate that the RSC flexibly constructs axis- and geometry-based reference frames, linking environmental structure to behaviorally relevant spatial codes. Our results extend these findings to freely moving humans using noninvasive recordings, showing that even without external visual landmarks, a comparable axis-aligned pattern emerges, anchored to the body’s principal axes.

To conclude, we provide compelling evidence in freely moving humans that theta activity arising from the RSC supports anchoring of space to the body’s principal axes. The effect was present only during goal-directed rotations, specific to the theta band and correlated with pointing performance, implying a self-motion–gated, axis-aligned compass in the RSC. Leveraging wearable sensors and advanced signal processing to examine brain activity during active movement in natural settings, these findings unify evidence from rodent and intracranial studies (see also Griffiths et al., 2024) and open the door to a new, ecologically grounded human electrophysiology framework.

## Method

In this analysis, we focused specifically on the rotation phases from a previous experiment (Saulay-Carret et al., 2025). These rotation data were not included in the analyses of the original publication.

### Experimental procedure

We recorded the EEG activity of 40 participants, and after exclusion of 2 due to motion sickness 38 were retained for the final analyses (*M =* 23.63 years; *SE* = 3.59 years; range 18-35 years; 22 female participants). Participants were immersed in a virtual arena with no external directional cues and asked to perform an adapted and extended version of the immersive viewpoint transformation task (He et al., 2022). They first performed a perspective taking task on a 2D map (for details see Saulay-Carret *et al*., 2025), in which they had to imagine being oriented in a certain direction and to retrieve the angle toward a target. Then they physically had to perform the rotations on themselves toward the indicated goal (referred to as goal-directed rotation hereafter), and we specifically analyzed these rotations. Once they were facing the intended direction, they validated their answer and performed the contrary rotation to return to the initial heading (referred as guided rotation, because four directional arrows appeared and guided participants to the starting direction) where a new map appeared after 500 ms. They completed 8 blocks of 62 trials each, for a total of 496 trials and approximately 75 minutes in the virtual environment. They performed rotations of 31 angles (every 12° between 0° and 360°), and each angle was repeated 9 times, alternating between left and right directions.

### Preprocessing and clustering of dipoles

EEG data were recorded at a 500 Hz sampling rate using a 64-channel Waveguard original cap with Ag/AgCl electrodes, referenced to CPz and grounded at AFz, with impedances kept below 10 kΩ. Signals were digitized using an eego mylab amplifier (24-bit resolution) and synchronized with stimuli via LabStreamingLayer (Kothe et al., 2025). Preprocessing was conducted offline in MATLAB (R2024a) using EEGLAB and the BeMobil pipeline (Klug et al., 2022). Data were first down-sampled to 250 Hz, and segments with inactivity or artifacts were removed. Spectral noise (*e*.*g*., line noise, screen refresh) was cleaned using Zapline-plus (Klug & Kloosterman, 2022), and noisy electrodes were identified and removed (on average, 2.18 ± 1.45 per participant), then reconstructed via spherical interpolation and re-referenced to the common average. A transient high-pass filter at 1.5 Hz was applied before running independent component analysis (ICA) with the AMICA (Klug & Gramann, 2021; Palmer et al., 2011) algorithm (2000 iterations, 10 rejections, 3 SD threshold). DipFit was used for dipole modeling, and components were classified using ICLabel (Pion-Tonachini et al., 2019). ICs identified as brain-related (≥ 30% probability, < 15% residual variance; Delorme et al., 2012) were retained (avg. 20 ± 3.96 per participant), and this information was transferred to the unfiltered cleaned data. The final signal was bandpass filtered (0.3–80 Hz), before applying source reconstruction focused on the retrosplenial complex (MNI: 0, -55, 15), using features from each IC (spectrum, ERSP, topography, dipole) to form 10-dimensional vectors. Clustering was done over 5000 iterations into 15 clusters, evaluated using six weighted metrics including participant representation in the cluster, residual variance, and distance to the region of interest (ROI). The final solution included 31 participants, with a mean residual variance of 0.056 and a mean distance of 16.03 mm from the ROI. For participants with multiple ICs in a cluster, we selected the closer IC based on the Euclidian distance from the ROI.

### Statistical analysis

To quantify oscillatory bursts in the dipoles clustered around the RSC, we applied the fBOSC framework (Seymour et al., 2022) to single-trial IC activations obtained from the broadband-filtered EEG signal (0.5-50Hz). Methodologically, fBOSC extends the Better OSCillation detection (BOSC) burst-detection framework by replacing BOSC’s background fit with Fitting Oscillations & One-Over-F (FOOOF; Donoghue et al., 2020) spectral parameterization, including a 1/f knee before applying BOSC’s power and duration-threshold criteria, thereby standardizing burst sensitivity across frequencies and improving robustness to spectral peaks. Movement-aligned rotation epochs were reconstructed from head-tracking data and resampled to match EEG sampling resolution. Trials were trimmed to exclude initial movement angles < 3° and retained only if the final rotation exceeded 90°, ensuring consistent movement engagement. Epochs were further screened using a robust quality-control pipeline excluding outliers based on amplitude, kurtosis, flat segments, and minimum duration criteria (on average 179.15 ± 23.29 epochs retained for the goal-direction rotation and 181 ± 21.87 for the guided rotation).

For each subject, fBOSC was run on the cleaned IC time courses using a Morlet wavelet decomposition (2–40□Hz; wavenumber = 5), with aperiodic background (1/f) estimation via FOOOF in knee mode (up to 4 peaks). Oscillatory episodes were identified where power exceeded the 95th percentile of the background-corrected power spectrum for at least two cycles at the given frequency. The resulting episode tables included the frequency, onset time, and duration for each burst. To test directional anchoring of theta episode onsets, we aligned each episode onset with the corresponding facing angle.

To quantify directional bias, we binned theta onset directions into 3° bins, excluding the first bin of each side, to ensure that we analyzed only the rotation periods, and computed the probability distribution of onset directions. A GLME with a Poisson distribution and log link was then fitted to the binned episode counts, with subject as random intercept. The model included circular predictors (cos□θ, sin□θ) to account for general periodic trends and two Gaussian-shaped anchoring regressors (σ□=□30°) centered at 90° and 270°, capturing directional clustering near cardinal axes. An interaction term between the two anchor regressors was included to account for joint effects. Model performance was assessed via likelihood-ratio tests comparing the full model (including anchor terms) against a reduced circular-only model. Improvements in fit were evaluated using deviance reduction and χ^2^ statistics. Statistical significance of individual predictors was assessed via Wald *z*-statistics, derived from the estimated coefficient divided by its standard error. The model also generated predicted onset probabilities, which were compared to observed probabilities across angles. Finally, we plotted the mean speed and acceleration at each directional bin to characterize movement dynamics across heading directions. For the correlational analyses, *β* estimates from the GLME at 90° and 270° were combined into a subject-level anchoring index and z-scored across participants. We then assessed the relationship of the z-scored values to mean pointing performance using both Pearson and Spearman correlations.

## Author Contributions

**Clément Naveilhan**: Conceptualization; Formal analysis; Investigation; Methodology; Writing—Original draft; Writing—Review & editing.

**Stephen Ramanoël**: Conceptualization; Funding Acquisition; Methodology; Project administration; Supervision, Writing—Review & editing.

## Acknowledgements

The authors acknowledge the essential contribution of the volunteer participants to the successful completion of this research. We also thank Maud Saulay-Carret for her help during data acquisition and thank Catherine Buchanan for her careful reading of the manuscript and her feedback.

This work was supported by the French government through the France 2030 investment plan managed by the National Research Agency (ANR), as part of the Initiative of Excellence Université Côte d’Azur under reference number ANR-15-IDEX-01 and, in particular, by the interdisciplinary Institute for Modeling in Neuroscience and Cognition (NeuroMod) of Université Côte d’Azur.

## Data and code availability

All the raw data, the analysis codes and the virtual environment generated for the present study are available online on the OSF repository of the study : https://osf.io/7bzua/?view_only=c087063470cb4cd18682ae83767d339f.

## Notes

### Competing Interest Statement

The authors have declared no competing interest.

